# Red fox viromes across an urban-rural gradient

**DOI:** 10.1101/2020.06.15.153858

**Authors:** Sarah J Campbell, Wilbur Ashley, Margarita Gil-Fernandez, Thomas M. Newsome, Francesca Di Giallonardo, Ayda Susana Ortiz-Baez, Jackie E Mahar, Alison L Towerton, Michael Gillings, Edward C Holmes, Alexandra JR Carthey, Jemma L Geoghegan

**Affiliations:** Department of Biological Sciences, Macquarie University, Sydney, NSW 2109, Australia; School of Life and Environmental Sciences, The University of Sydney, Sydney, NSW 2006, Australia; Marie Bashir Institute for Infectious Diseases and Biosecurity, School of Life and Environmental Sciences and School of Medical Sciences, The University of Sydney, Sydney, NSW 2006, Australia; The Kirby Institute, University of New South Wales, Sydney, NSW 2052, Australia; Greater Sydney Local Land Services, Sydney, NSW, Australia; Department of Microbiology and Immunology, University of Otago, Dunedin 9016, New Zealand; Institute of Environmental Science and Research, Wellington 5018, New Zealand

**Keywords:** *Vulpes vulpes*, carnivore, predator, canine, exotic species, urban, virus, metagenomics

## Abstract

The Red fox (*Vulpes vulpes*) has established large populations in Australia’s urban and rural areas since its introduction following European settlement. Foxes’ cryptic and highly adaptable nature allows them to invade cities and live among humans while remaining largely unnoticed. Urban living and access to anthropogenic food resources also influences fox ecology. Urban foxes grow larger, live at higher densities and are more social than their rural counterparts. These ecological changes in urban red foxes are likely to impact the pathogens that they harbour, and foxes could pose a disease risk to humans and other species that share these urban spaces. To assess this possibility, we used a meta-transcriptomic approach to characterise the viromes of urban and rural foxes across the Greater Sydney region in Australia. Urban and rural foxes differed significantly in virome composition, with rural foxes harbouring a greater abundance of viruses compared to their urban counterparts. In contrast, urban fox viromes comprised a greater diversity of viruses compared to rural foxes. We identified nine potentially novel vertebrate-associated viruses in both urban and rural foxes, some of which are related to viruses associated with disease in domestic species and humans. These included members of the *Astroviridae, Picobirnaviridae, Hepeviridae* and *Picornaviridae* as well as rabbit haemorrhagic disease virus-2 (RHDV2). This study sheds light on the viruses carried by urban and rural foxes and emphasises the need for greater genomic surveillance of foxes and other invasive species at the human-wildlife interface.

**Importance:** Urbanisation of wild environments is increasing as human populations continue to expand. Remnant pockets of natural environments and other green spaces in urban landscapes provide invasive wildlife such as red foxes with refuges within urban areas, where they thrive on the food resources provisioned by humans. Close contact between humans, domestic species and foxes likely increases the risk of novel pathogen emergence. Indeed, the vast majority of emerging infectious diseases in humans originate in wild animals. Here, we explored potential differences in viromes between urban fox invaders and their rural counterparts. Viromes of foxes and their ectoparasites comprise a diversity of viruses including those from the *Astroviridae, Picobirnaviridae, Hepeviridae, Caliciviridae* and *Picornaviridae*. Microbial surveillance in foxes and other urban wildlife is vital for monitoring viral emergence and for the prevention of infectious diseases.

## Introduction

Red foxes (*Vulpes vulpes*) have the largest natural distribution of any wild terrestrial carnivore (1), extending through Eurasia and north America (2). After their introduction into Australia in the mid-1800’s, their range expanded to cover most of the continent. Red foxes exploit a wide range of habitats with varying climates, from alpine to desert, and are considered one of the most adaptable species on the planet. They are broadly distributed across different land uses including natural and forested landscapes as well as highly urbanised, human dominated landscapes (3, 4). Red fox home range size varies depending on resource availability and land use type. Globally, urban fox home ranges average approximately 1.7 km^2^, while rural fox home ranges are larger, at around 5.7 km^2^ on average (5). In Australia, home ranges for foxes in arid regions can reach at least 120 km^2^ (6), between 5-7km^2^ in rural areas (7) and less than 1km^2^ in urban centres (8).

Foxes are common across rural and bushland regions in Australia and have established a large presence in major metropolitan centres (3, 9), being recorded near the Sydney region since 1907 (10). They were first sighted in an Australian city (Melbourne) in 1943, although they were noted in Melbourne’s suburban surrounds as early as 1933 (11). For comparison, foxes were first noted in British cities (i.e. in their native range) in 1930 (4). Urban cities support much higher densities of foxes than more rural regions. In Melbourne, city foxes live in densities of up to 16 individuals per km^2^ (9). This is compared to just 0.2 individuals per km^2^ in more rural areas (3). In Bristol city in the UK, densities as high as 35 individual foxes per km^2^ have been estimated (12).

Predation by red foxes is a key threat to Australian native fauna (13). The list of native animals threatened by fox predation includes some of Australia’s most endangered species, such as the rufous hare-wallaby (*Lagorchestes hirsutus*) and loggerhead turtle (*Caretta caretta*), as well as the critically endangered brush-tailed bettong (*Bettongia penicillata*), Gilbert’s potoroo (*Potorous gilbertii*), western swamp tortoise (*Pseudemydura umbrina*) and orange bellied parrot (*Neophema chrysogaster*) (14). Due to the threat foxes pose to endangered wildlife and Australian biodiversity, fox populations are actively controlled. Poison baiting is the most common and cost-effective method of control in rural areas (3). In urban areas, however, the risk to pets limits control methods to trapping and shooting (15). However, foxes are notoriously difficult to trap and shooting in urban areas requires tracking at night (when foxes are most active) by licensed professionals. Such limitations make it difficult to effectively control foxes in urban areas.

Red foxes exhibit cryptic and nocturnal behaviour, going largely undetected in urban areas despite their high abundance (16, 17). They thrive on the resources inadvertently provided by humans in cities and may develop distinct urban behaviours as a consequence of urban living (4, 18, 19). For example, urban carnivores such as coyotes (*Canis latrans)* display increased boldness and decreased human aversion when compared to their rural counterparts (4, 20, 21). Urban living also increases carnivore body size which may have positive effects on fitness and fecundity (4, 22). When food is abundant, carnivore home ranges are smaller, higher densities are supported and encounters between conspecifics are more frequent (4, 23, 24). In urban areas, fox family group sizes are often larger than those in rural areas, with juvenile females remaining in their natal territory to assist with cub rearing (9, 25, 26). Thus, urban environments may favour increased conspecific tolerance and social behaviours in foxes (9, 24-26).

Although red foxes are known to harbour a diversity of viruses (27, 28), it is unknown whether urban and rural foxes have different viral compositions. High-density living and increased host contact can increase pathogen transmission rates among hosts (29). As such, a high-density population of cryptic urban foxes living in close proximity to largely unsuspecting humans could pose an important pathogen risk. Foxes are likely to investigate human refuse, including compost and rubbish bins, and consume food scraps from surfaces such as outdoor barbeques and furniture, eat from pet bowls and wildlife feeding stations and defecate nearby, increasing the potential for pathogen transfer (18). In addition, as urban animals can gradually become habituated to humans (4), we would expect to see an increase in direct fox-human interactions with the potential for disease transmission between the two species.

Using an unbiased, meta-transcriptomic approach, we describe, for the first time, the virome of the introduced Australian red fox sampled from urban and rural regions. We hypothesised that foxes in urban areas could harbour a greater viral diversity and abundance compared to rural foxes, due to the potentially higher population densities and increased conspecific interactions in urban areas. While there is limited information on fox social dynamics in Australia, we also postulated that females could harbour a greater diversity and abundance of viruses than males due to particular social behaviours reported for female foxes in their native ranges, such as cooperative cub rearing (25, 26). To this end, samples (liver, faecal and ectoparasite) were collected from foxes around the Greater Sydney region, Australia, including in urban and more rural areas (Figure 1). Samples were pooled (based on sampling location and sex) and subject to RNA sequencing to reveal viral diversity, evolution and abundance.

**Figure 1.**
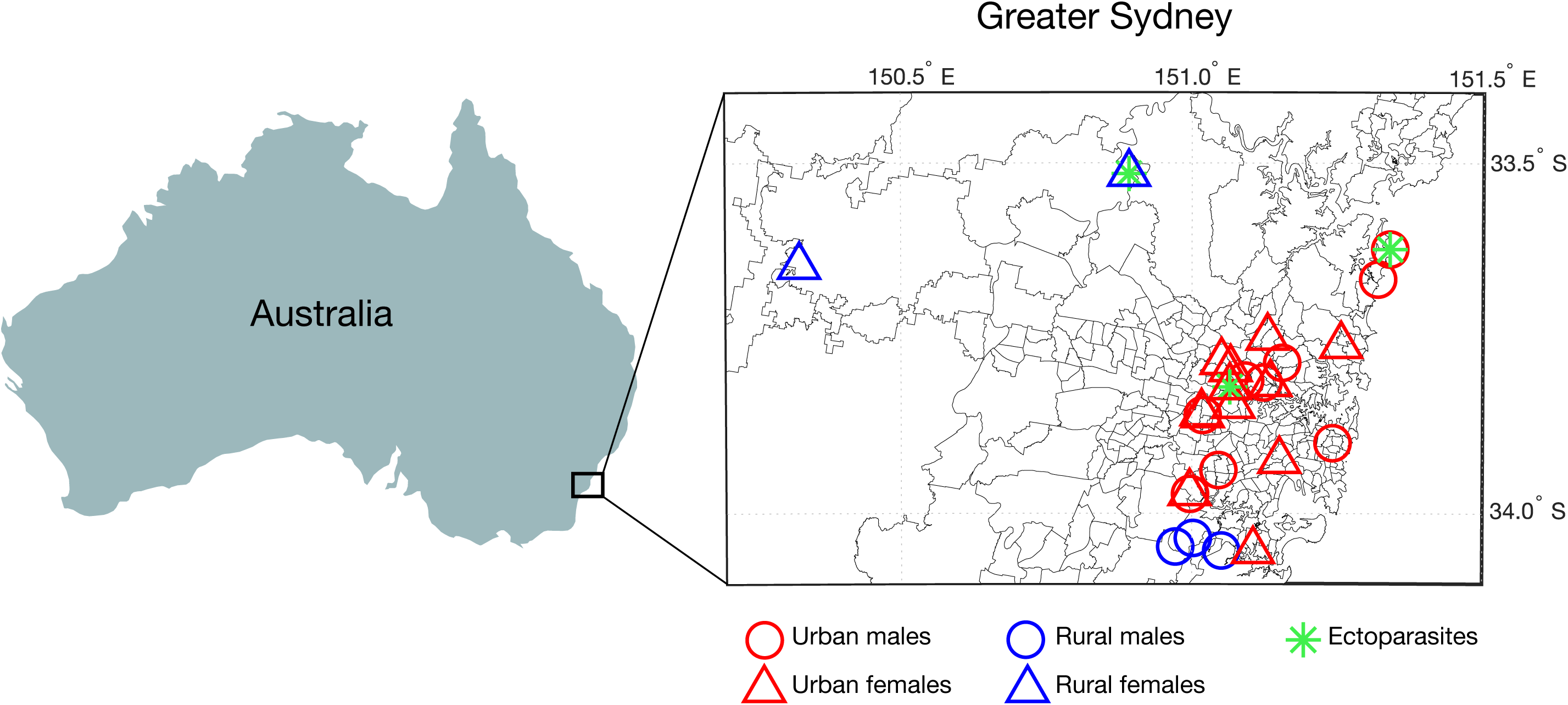
Map of the Greater Sydney region showing fox sampling locations of urban (red) and rural (blue) fox carcasses, identified as male (circle) or female (triangle), as well as those harbouring ectoparasites (green asterisk).

## Materials & Methods

### Sample collection

The current project was part of a larger research program into urban foxes in partnership with Greater Sydney Local Land Services, a New South Wales State Government organisation responsible for management of pest species across the region. We collected fresh carcasses from independent licensed trappers and shooters who were actively controlling foxes in the Greater Sydney region (see Figure 1 for sample locations). To minimise degradation of RNA, samples were taken as soon as possible after death (03:19:00 ± 02:59:00 hrs post-mortem, n=27). One carcass had been frozen for approximately one week and one carcass had been dead for an unknown amount of time. The foxes used for this study were either trapped in cages and shot, or tracked and shot. One individual was obtained as recent roadkill. Foxes killed by poison baits were excluded.

Whole fox carcasses were collected and transported to the laboratory where they were immediately dissected to collect faecal, liver and ectoparasite samples. All samples were individually stored in RNALater at −80°C. We sampled a total of 29 individual foxes; 13 males and 16 females. For this study, foxes were classified as juvenile if their body mass and body length were less than 3.3 kg and 51cm, respectively. These values were chosen as the body mass of an adult red fox can range between 3.3 and 8.2kg, while body length can range between 51 and 78cm (when measured from the tip of the nose to the first vertebra of the tail) (30). Based on this assessment, 25 foxes were classified as adults (12 males, 13 females) and four as juveniles (1 male, 3 females).

### Sampling in urban and rural areas

Fox sampling relied on coordination with professional pest control operators who focus control efforts in specific locations in accordance with local control initiatives. For this reason, a representative sample across a land-use gradient from urban to rural was not possible. Sufficiently fresh rural and bushland fox samples were also difficult to obtain since poison baiting is the principal control method in these areas. Therefore, ‘rural’ was broadly defined as any natural bushland, national park, mostly agricultural or sparsely populated region outside the central urban districts, with a human population density of fewer than 500 people per km^2^. Similarly, ‘urban’ was defined as built up areas inside the central urban district (including parks, gardens and golf courses) with a population density of more than 500 people per km^2^ either in the area sampled or in the immediate surrounding areas. Human population density information was obtained from the Australian Bureau of Statistics (2016 census data) (31). Central urban districts were defined by the Urban Centres and Localities statistical classification (UCL) (32). Land use classification and human population density cut-offs were loosely based on work by Stepkovitch (2019).

### RNA extraction and whole transcriptome sequencing

Qiagen RNeasy Plus Mini Kits were used to extract RNA from liver, faecal, and ectoparasite samples from collected red fox carcasses. Thawed samples were transferred to a lysis buffer solution containing 1% β-mercaptoethanol and 0.5% Reagent DX. Samples were homogenised and centrifuged. DNA was removed from the supernatant via gDNA eliminator spin column and RNA was eluted via RNeasy spin column. RNA concentration and purity were measured using the Thermo Fisher Nanodrop. Samples were pooled based on land use category (urban or rural), sex and sample type (liver, faecal or ectoparasite), resulting in nine representative sample pools (Table 1). Adults and juveniles were pooled as only two juveniles were sampled. Ectoparasites included fleas (*Siphonaptera*) and ticks (*Ixodida*). These were not classified below the Order level and due to the small number sampled were also pooled. The TruSeq Stranded Total RNA Ribo-Zero Gold (h/m/r) kit was used to prepare pooled samples for sequencing. Pooled samples were sequenced on the NextSeq 500 with 2×75bp output at the Ramaciotti Centre for Genomics at the University of New South Wales, Sydney. Sequencing resulted in nine representative data libraries (Table 1). The raw reads are available on NCBI’s SRA database under BioProject XXX, while the consensus sequences of each virus have been submitted to NCBI GenBank and assigned accession numbers XXX-YYY.

**Table 1.**
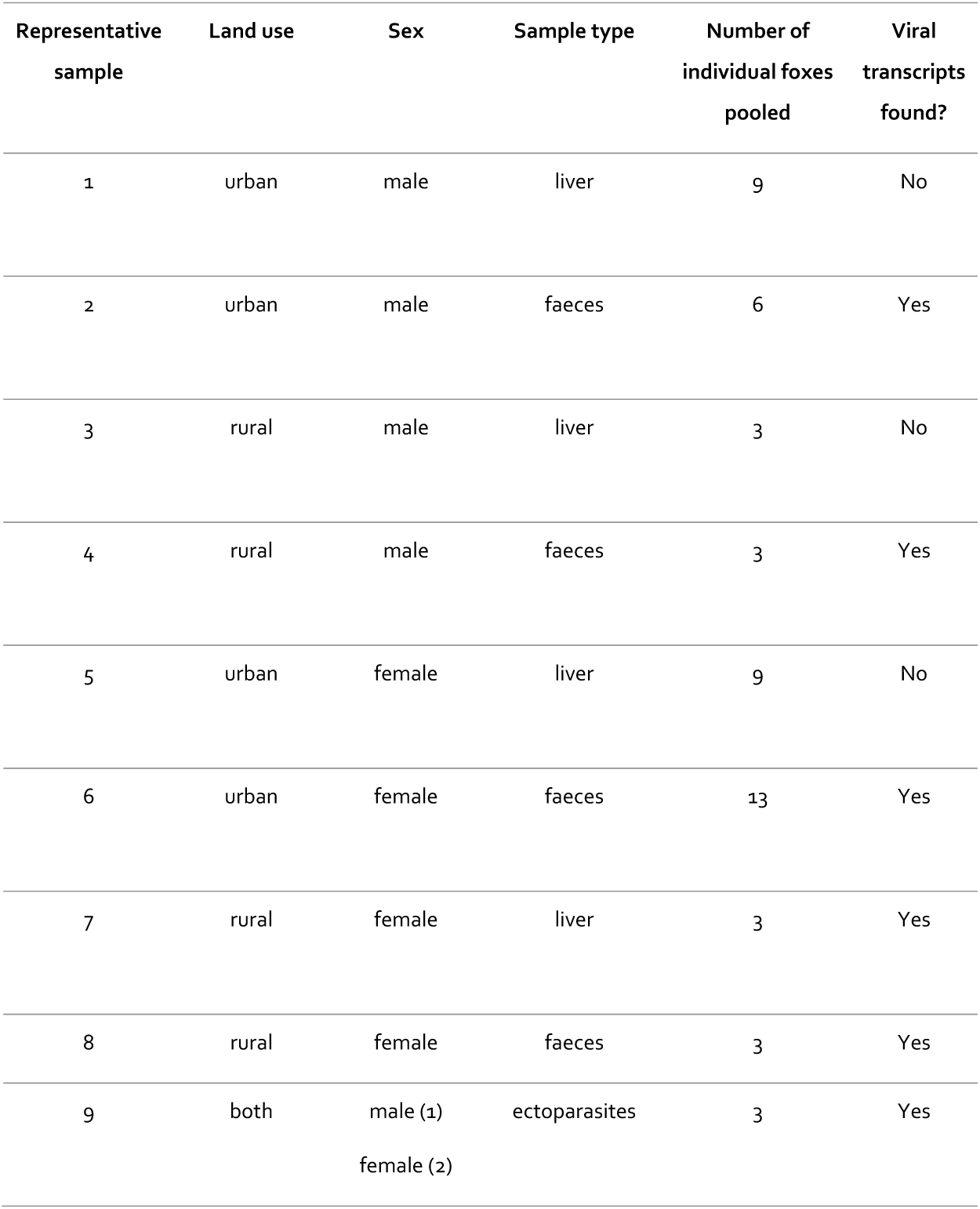
Breakdown of red fox representative samples, detailing land-use, sex and sample type, as well as the number of individuals pooled for RNA sequencing.

| Representative sample | Land use | Sex | Sample type | Number of individual foxes pooled | Viral transcripts found? |
| --- | --- | --- | --- | --- | --- |
| 1 | urban | male | liver | 9 | No |
| 2 | urban | male | faeces | 6 | Yes |
| 3 | rural | male | liver | 3 | No |
| 4 | rural | male | faeces | 3 | Yes |
| 5 | urban | female | liver | 9 | No |
| 6 | urban | female | faeces | 13 | Yes |
| 7 | rural | female | liver | 3 | Yes |
| 8 | rural | female | faeces | 3 | Yes |
| 9 | both | male (1)<br>female (2) | ectoparasites | 3 | Yes |

### Virus discovery

Sequencing reads were assembled *de novo* into longer sequences (contigs) based on overlapping nucleotide regions using Trinity RNA-Seq (33). Assembled contigs were assigned to a taxonomic group (virus, Bacteria, Archaea, Eukarya) and viruses were identified to their closest species match based on sequence similarity searches against the NCBI nucleotide (nt) and non-redundant protein (nr) databases using BLASTn (34) and Diamond (BLASTX) (35), respectively. An e-value threshold of 1×10^−5^ was used as a cut-off to identify positive matches. We removed non-viral hits, including host contigs with similarity to viral sequences (e.g. endogenous viral elements), as well as any contigs with similarity to plant viruses, which were more likely to be derived from the foxes’ diet.

### Inferring the evolutionary history of fox viruses

We inferred the phylogenetic relationships of the vertebrate-associated viruses identified in the fox samples. Vertebrate-associated viruses were defined as viruses which shared sequence similarity to other known vertebrate viruses. First, the amino acid translations of the viral transcripts were combined with other virus protein sequences from the same virus families obtained from GenBank (Table 2). Second, the sequences were aligned using MAFFT v.3.4, employing the E-INS-I algorithm. Ambiguously aligned regions were removed using trimAl v.1.2 (36). To estimate phylogenetic trees, we selected the optimal model of amino acid substitution identified using the Bayesian Information Criterion as implemented in Modelgenerator v0.85 (37) and employed the maximum likelihood approach available in PhyML v3.1 (38) with 1000 bootstrap replicates. For the viral transcript matching RHDV2 we used a nucleotide alignment with similar viruses. New viruses were named after fictional fox characters.

**Table 2.** Vertebrate-associated viral contigs, contig length (nt), percent abundance in their respective pools and the percent amino acid identity to their closest match on NCBI/GenBank.

| Land use (sex) | Virus name (species) | Virus family | Contig length (nt) | % Relative abundance | Closest match (GenBank accession number) | % Amino acid identity |
| --- | --- | --- | --- | --- | --- | --- |
| rural (female) | Vixey virus | *Picornaviridae* | 2427 | 0.007% | Canine kobuvirus (AZS64124.1) | 97.65% |
|  | Wilde virus-1 | *Picornaviridae* | 7236 | 5.66% | Canine picornavirus (YP_005351240.) | 89.18% |
|  | Swiper virus | *Hepeviridae* | 7374 | 0.01% | Elicom virus-1 (YP_009553584.) | 28.92% |
|  | Red fox associated rabbit haemorrhagic disease virus-2 | *Caliciviridae* | 7026 | 0.14% | Rabbit haemorrhagic disease virus-2 (MF421679.1) | 99.62% |
| rural (male) | Tod virus-2 | *Picornaviridae* | 4263 | 0.17% | Canine picodistovirus (YP_007947664.) | 98.53% |
|  | Vulpix virus | *Astroviridae* | 2556 | 0.046% | Feline astrovirus (YP_009052460.) | 96.11% |
| urban (female) | Tod virus-1 | *Picornaviridae* | 2062 | 0.0004% | Canine picodistovirus (YP_007947664.) | 98.83% |
|  | Charmer virus | *Picobirnaviridae* | 448 | 0.0001% | Wolf picobirnavirus (ANS53886.1) | 80.27% |
| urban (male) | Wilde virus-2 | *Picornaviridae* | 1524 | 0.00058% | Canine picornavirus (YP_005351240.) | 73.37% |

### Diversity and abundance analysis

Transcript abundance for all viruses (vertebrate and invertebrate-associated) was estimated using RSEM within Trinity (39). Specifically, we assessed how many short reads within a given library mapped to a particular transcript. Raw counts were then standardised against the total number of reads within each library. Virome diversity (i.e. virus species richness) and relative abundance were compared among samples using a non-metric multidimensional scaling (nMDS) ordination in conjunction with an analysis of similarities (ANOSIM) based on Bray-Curtis dissimilarity as implemented in the vegan package in R (40). To determine which viral families were contributing the most to differences between samples, an indicator species analysis was performed, using a point biserial coefficient of correlation within the indicspecies package in R (41).

## Results

Meta-transcriptomic sequencing of nine representative pooled samples resulted in 44-57 million paired reads per pool (593,406,706 reads in total). BLAST analyses revealed that the faecal samples were dominated by bacteria (51.17-84.61%), while the liver samples were dominated by eukaryotic transcripts (92.90-99.43%), largely comprising fox RNA. Viruses made up a small proportion of the four representative faecal samples (0.002-5.85%) and were detected in only one of the representative liver samples (0.001%). Archaea were detected at very low levels in faecal samples only (0.002-0.021%). The ectoparasites (fleas and ticks) differed substantially to the liver and faecal samples with 50.97% of reads classed as ‘unmatched’ meaning they did not share sequence similarity to any known sequence. The remainder of the contigs from ectoparasite samples were from eukaryotes (44.39%), bacteria (4.64%) and viruses (0.004%). Unmatched reads in liver and faecal samples ranged between 0.52-12.22% (Figure 2a).

**Figure 2.**
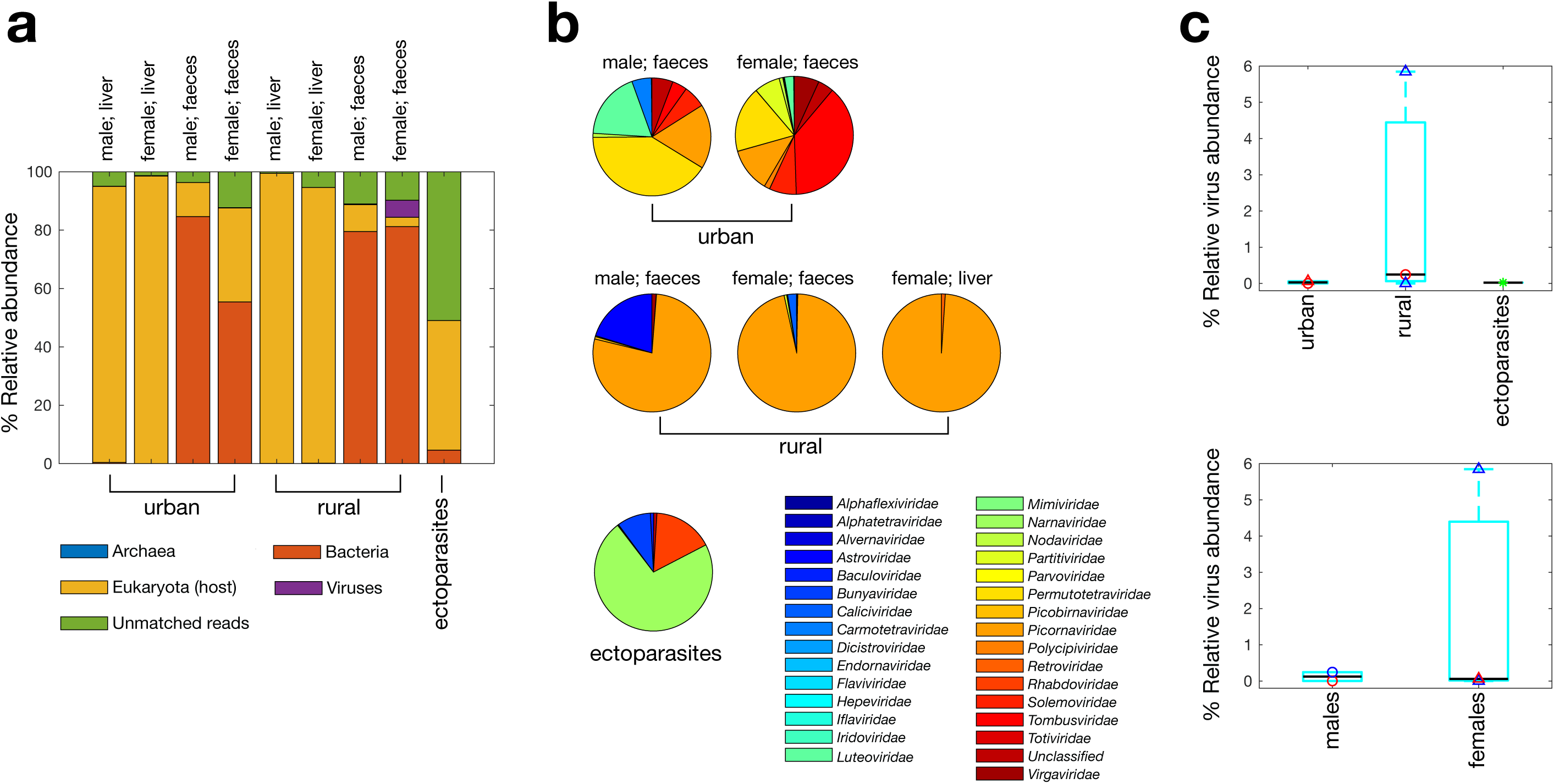
Overview of the red fox virome. (a) Percentage abundance of each taxonomic group identified in each respective pooled sample, standardised against the number of raw reads per pool. Due to their low abundance, archaea (0.002-0.021%) and some of the viral reads (0.001-5.85%) are too small to visualise. (b) Percentage abundance of (eukaryotic-associated) viral families detected in each respective pooled sample (excluding bacteriophage). (c) Boxplots showing percentage abundance of (eukaryotic-associated) viral reads in urban, rural and ectoparasite samples and males and females. A black line indicates the median and the bottom and top edges of the box indicate the 25th and 75th percentiles, respectively. Raw abundances are superimposed, and the colour and shape of data points are as in Figure 1.

Multiple novel vertebrate-associated virus transcripts were identified from both urban and rural foxes, including a hepevirus, picobirnavirus, astrovirus and various picornaviruses (Table 2). In addition, we found virus transcripts with sequence similarity to rabbit haemorrhagic disease virus-2 (RHDV2). Vertebrate-associated virus transcripts represented between 0.4-98% of viral reads. The remainder comprised mostly invertebrate, plant and fungi associated virus transcripts which were most likely acquired from the foxes’ diet.

### Virome composition

Urban, rural and ectoparasite samples had distinctly different virome compositions (ANOSIM R = 1, *p* = 0.0167; Figure 2 and Figure 3). Transcripts from a total of 30 distinct viral families were identified across the six pools in which viral RNA was detected (rural male faeces, rural female faeces, rural female liver, urban male faeces, urban female faeces, and ectoparasites). Overall, 21 viral families were identified in transcripts from urban foxes and 19 from rural foxes. Urban foxes exhibited a higher diversity of viruses compared to rural foxes; transcripts from the latter were heavily dominated by *Picornaviridae*, which made up between 77.33-98.97% of the virome of rural foxes (Figure 2b). Indicator species analysis suggested that while the rural samples were characterised by the presence of *Picornaviridae* (stat = 0.978, *p* = 0.0496), the urban samples were significantly associated with the presence of *Nodaviridae* (stat = 0.998, *p* = 0.0498). Viral diversity was higher in females (25 distinct viral families) than in males (13 distinct viral families). A much larger percentage of the viral transcripts identified were vertebrate-associated in rural foxes (male: 98.23%, female: 97.84%) compared to urban foxes (male: 2.41%, female: 0.39%), although this percentage was higher in males in both groups. In this context it is important to note that some virus transcripts found here may be the result of contamination by reagents.

**Figure 3.**
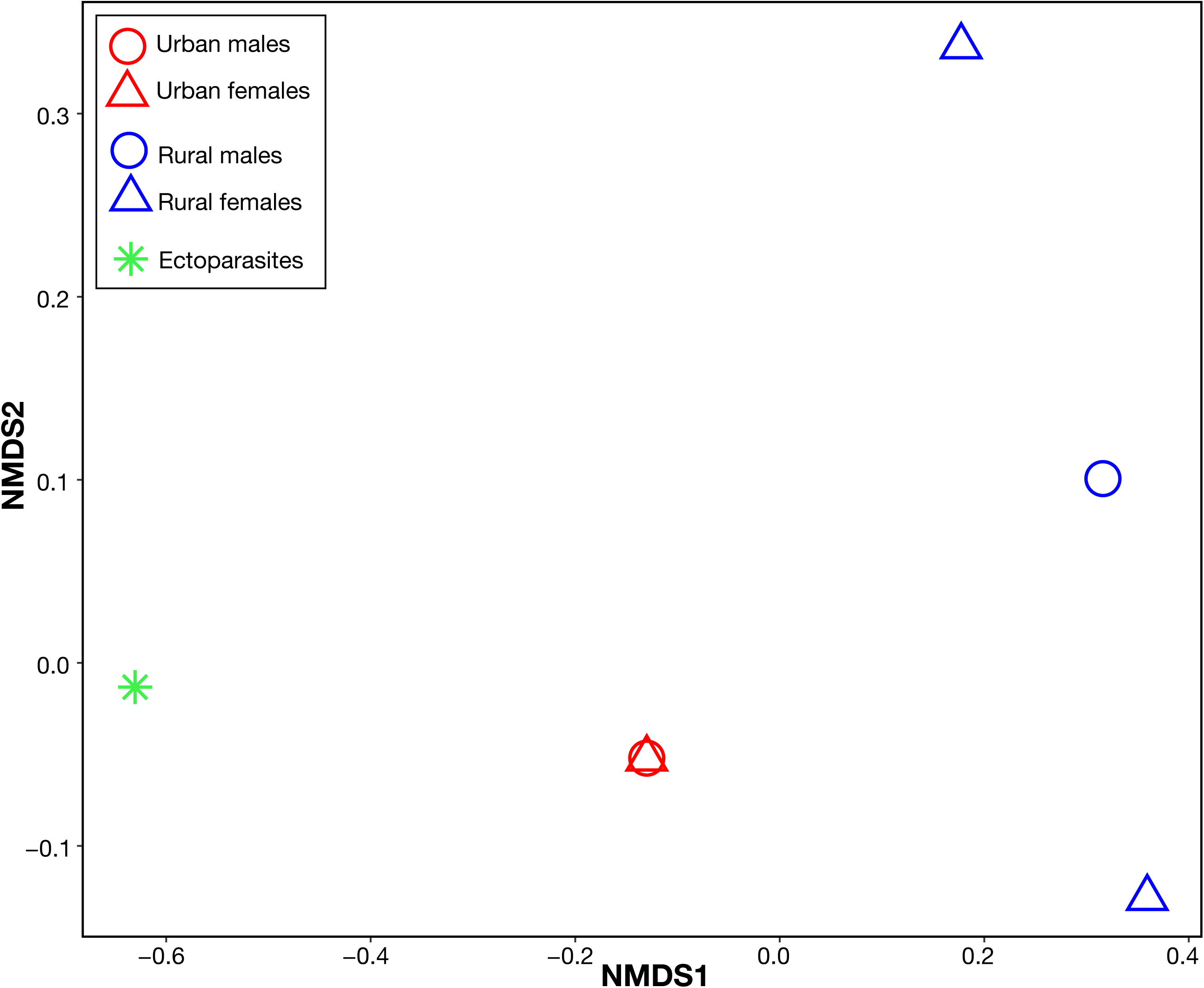
Non-metric multidimensional scaling (nMDS) ordination showing differences in virome composition (at the family level) among samples according to habitat and sex. Individual points represent individual pooled samples. Points closer together have a more similar virome composition (based on Bray-Curtis dissimilarity, which incorporates both the diversity and abundance of viruses) and *vice versa* for those further apart.

On average, total viral abundance (including both vertebrate and non-vertebrate viruses) was higher in rural foxes (2.03 ± 3.31%, n=3) than in urban foxes (0.03 ± 0.04%, n=2), and in female foxes (1.97 ± 3.36%, n=3) than in male foxes (0.12 ± 0.17%, n=2) (Figure 2c). However, due to the small sample size, differences between males and females may be due to individual animals contributing more to overall abundance than others. For example, the rural female fox pool (comprising three individual foxes) contained an unusually high number of viruses (>5%) compared to the others. This may have inflated virus abundance counts in females when combined. While virome composition was compared among a relatively small number of samples, this is balanced by the fact that each sample comprises the viromes of multiple individual foxes (n = 3-13 foxes per pool; Table 1).

### Vertebrate-associated viruses in foxes

#### Hepeviridae

Hepevirus (positive-sense single stranded RNA viruses) sequences were discovered in the rural female faecal samples. Tentatively named swiper virus, this virus transcript was very distinct in sequence, sharing only 28.92% amino acid identity to its closest relative, elicom virus-1 from mussels, and had a relative abundance of 0.01% (Table 2). While its closest genetic relative is not from a vertebrate host suggesting it may be a diet associated contaminant, phylogenetic analysis of the RNA dependant RNA polymerase (RdRp) encoding region placed this hepevirus in close proximity to both house mouse hepevirus and elicom virus-1, with these viruses forming a distinct monophyletic group (Figure 4).

**Figure 4.**
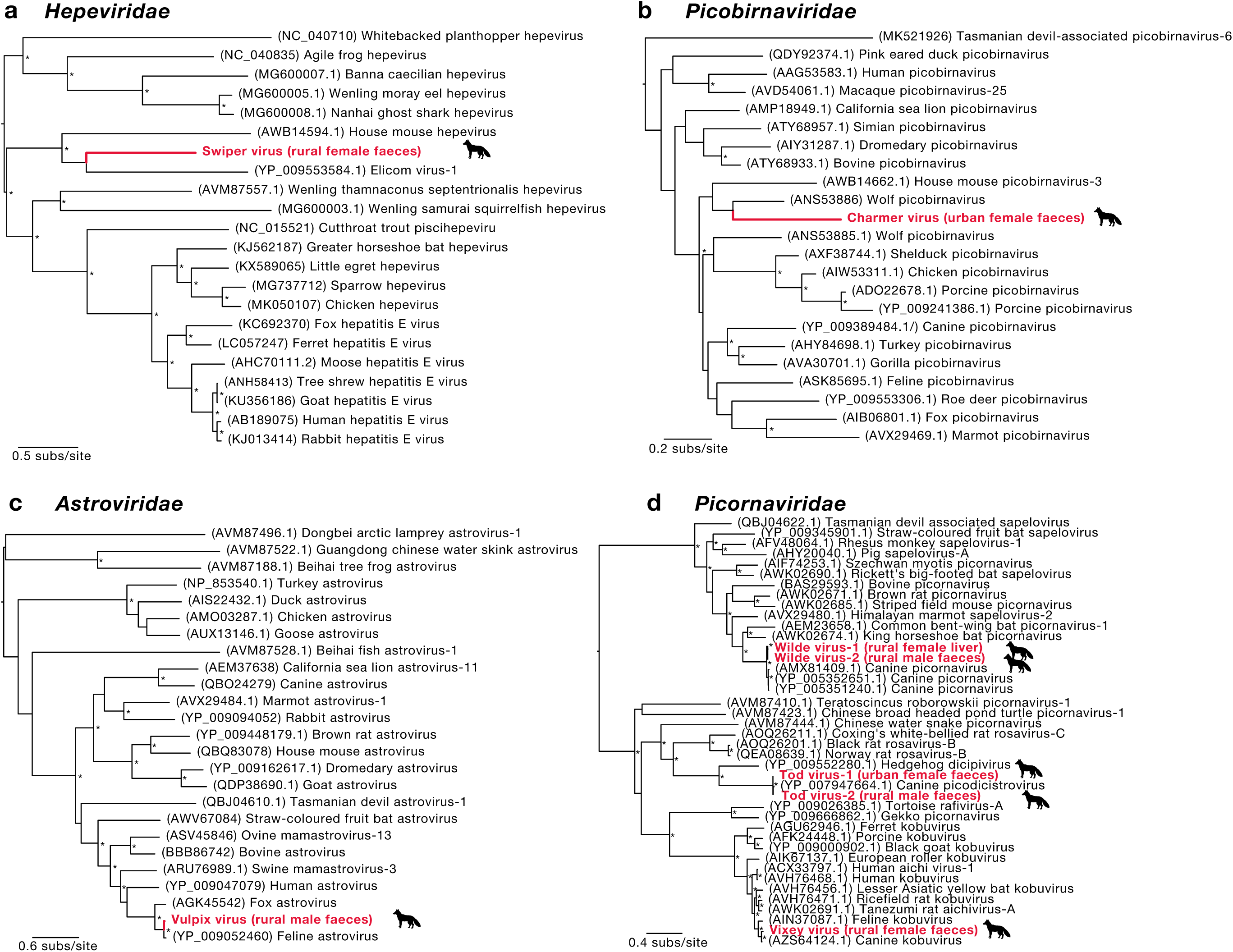
Phylogenetic relationships of likely vertebrate-associated viruses discovered from assembled contigs: (a) *Hepeviridae*, (b) *Picobirnaviridae*, (c) *Astroviridae* and (d) *Picornaviridae*. The maximum likelihood phylogenetic trees show the topological position of the newly discovered potential viruses (bold, red text), in the context of their closest relatives. All branches are scaled to the number of amino acid substitutions per site and trees were mid-point rooted for clarity only. An asterisk indicates node support of >70% bootstrap support.

#### Astroviridae

We detected an astrovirus (positive-sense single stranded RNA virus), tentatively named vulpix virus, in the rural male faecal samples. Notably, the sequence shared a 96.11% amino acid identity with feline astrovirus D1 and had a relative abundance of 0.046% (Table 2). Based on phylogenetic analysis of the RdRp, this virus clustered with other mammalian-associated viruses within the mamastroviruses (Figure 4).

#### Picobirnaviridae

Picobirnavirus (double-stranded RNA viruses) sequences were detected in urban male, rural male and urban female faecal samples. As some of the sequences represented less conserved regions of the viral genome, only one RdRp sequence (from the urban female samples) was used for phylogenetic analysis. The sequence, tentatively named charmer virus, shared an 80.27% amino acid identity with a picobirnavirus found in wolves and had a relative abundance of 0.0001% (Table 2). The sequence also clustered with other mammalian associated picobirnaviruses (Figure 4).

#### Picornaviridae

Several picornaviruses (positive-sense single stranded RNA viruses) were discovered. Two kobuvirus related sequences were discovered in the rural female faecal samples. The longer sequence, tentatively named vixey virus, shared highest amino acid identity with canine kobuvirus from a domestic dog (97.65%) and had a relative abundance of 0.007% (Table 2). Analysis of the RdRp region showed the sequence clustered most closely with feline kobuvirus and other mammalian kobuviruses (Figure 4).

Multiple picodicistrovirus sequences were detected in the urban male, rural male and urban female faecal samples. Two of the sequences, tentatively named tod virus-1 and tod virus-2 both shared 98% amino acid identity with canine picodicistrovirus (Table 2). Based on analysis of the RdRp region the sequences clustered together with mammalian dicipivirus and rosaviruses as well as reptilian picornaviruses (Figure 4).

Multiple picornavirus sequences were identified in the rural male and rural female samples. Two sequences, tentatively named wilde virus-1 and 2, all shared between 73-89% amino acid identity with canine picornavirus and had relative abundances of 5.66% and 0.00058%, respectively (Table 2). These sequences clustered with other mammalian picornaviruses in the order *Sapelovirus* (Figure 4).

#### Caliciviridae

One of the most striking observations was the identification of rabbit haemorrhagic disease virus-2 (RHDV2) (positive-sense single-stranded RNA virus) in rural female and urban male faecal samples. The viral sequence in the rural female samples shared a 99.62% amino acid identity with RHDV2 isolated from rabbits between 2015-2016 and had a relative abundance of 0.14% (Table 2) (Figure 5). The viral sequence in the urban male samples was too short to enable phylogenetic analysis. This is the second time that RHDV2 has been found in non-rabbit hosts (42), presumably through rabbit consumption in this case.

**Figure 5.**
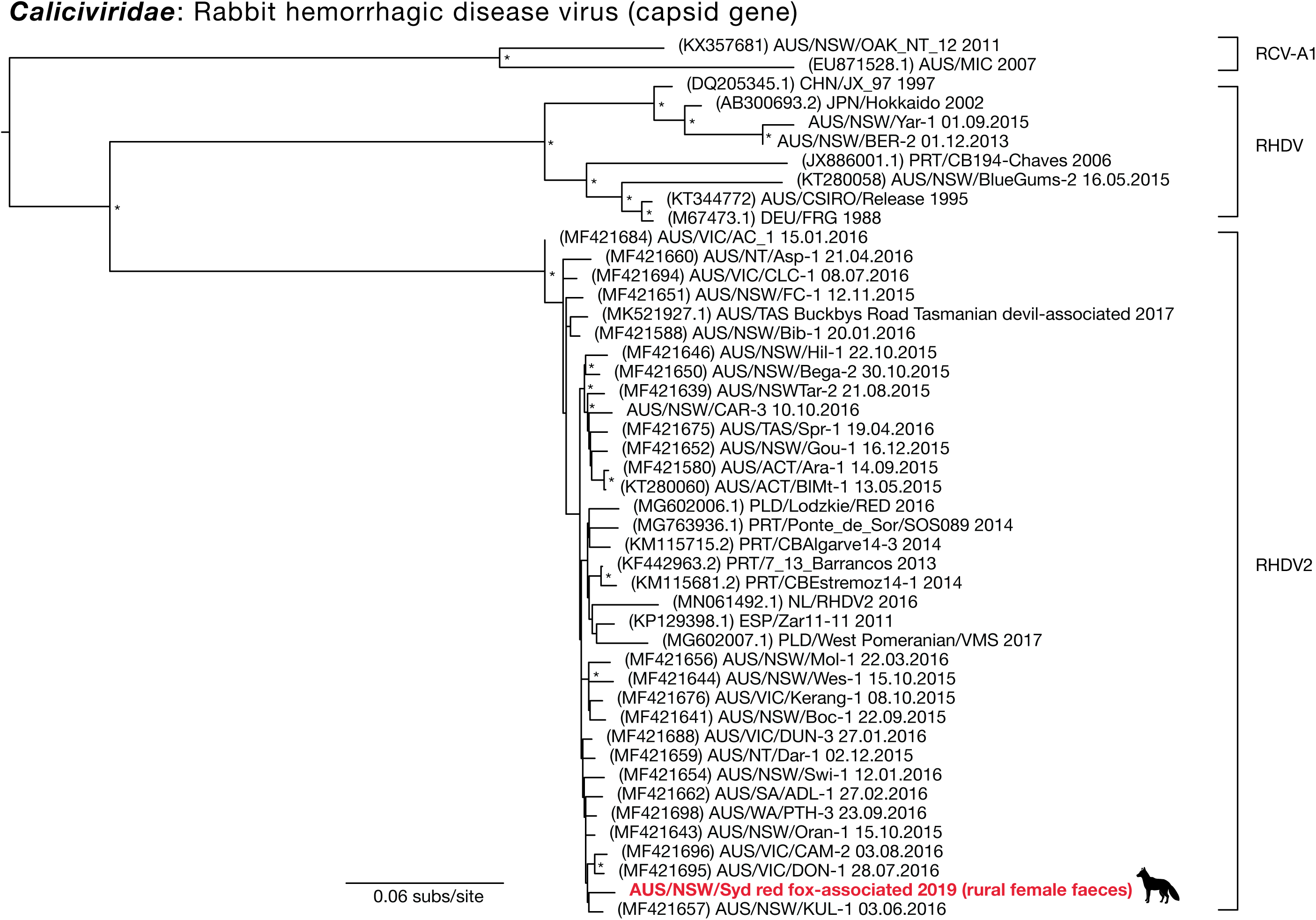
A maximum likelihood phylogenetic tree showing the topological position of RHDV2 capsid gene in the red fox (bold, red text), in the context of its closest relatives. Major clades are labelled. All branches are scaled to the number of nucleotide substitutions per site and trees were mid-point rooted for clarity only. An asterisk indicates node support of >70% bootstrap support.

## Discussion

We have revealed that Sydney’s red foxes, in both urban and rural environments, harbour a wide diversity of viruses, some of which are genetically similar to those that infect domestic pets and humans. Domestic mammals tend to hold central positions in mammal viral transmission networks (43). The close genetic similarity of the viruses found here to viruses frequently found in common domestic pets such as cats and dogs suggests cross-species transmission between foxes and domestic species may have occurred. The most cited case of viral transmission between humans and domestic pets is the transmission of rabies virus (44), although other examples include noroviruses from dogs, isolated cases of influenza A(H7N2) virus from cats (45, 46) and numerous bacterial diseases and parasites (44, 47). There may also be additional cases of viral sharing between humans and their pets, although these may go undiagnosed due to insufficient knowledge of the genetic variability of these viruses and their relationships with hosts.

All vertebrate-associated viruses found here were RNA viruses. Although this may in part be due to the reliance on transcript-based viral detection, RNA viruses are in general characterised by lower host specificity than DNA viruses, reflecting an increased occurrence of cross-species transmission (43, 48). The opportunity for interactions between urban wildlife, pets and humans provide likely transmission pathways for novel RNA viruses. Indeed, eukaryotic parasites are already known to infect human hosts following the wildlife-domestic pet-human transmission network (49). We discovered viral transcripts with some sequence similarity to the *Hepeviridae* that cause hepatitis E in mammals, which has already been isolated from various domestic and wild animals including foxes in the Netherlands (28, 50). Confirmed zoonotic cases include transmission to humans from domestic pigs, cats and wild rodents (50, 51). In contrast, the hepevirus detected here was phylogenetically distinct from the fox hepatitis E virus previously detected (28), and instead was more closely related to hepeviruses detected in freshwater mussels and a house mouse. Hence, although we have classed the virus as vertebrate-associated, its divergent phylogenetic position could in fact mean that it results from dietary consumption.

The astrovirus transcript (vulpix virus) showed the greatest sequence similarity (96%) to astroviruses from domestic cats as well as from other foxes, humans and pigs. Astroviruses have a broad host range (52) and are frequently detected in the faeces of mammals, birds and humans with gastroenteritis (53, 54). Astroviruses have also been associated with other diseases and disorders such as shaking syndrome in minks (55), neurological disease in cattle (56) and encephalitis in humans (57). Some human astroviruses are more closely related to those in animals than to each other, suggesting that these viruses periodically emerge from zoonotic origins (58). The similarity of fox astroviruses to those found in cats indicates that these viruses may have jumped hosts in the past and highlights further the potential role of domestic pets and wildlife in virus transmission.

Picobirnaviruses are found in humans and other mammals and are thought to be linked with gastroenteritis, however their role in disease remains unclear (59, 60). The picobirnavirus related transcript found here showed the greatest sequence similarly to a picobirnavirus found in wolves with diarrheic symptoms (59). It is also similar to picobirnaviruses described as potentially zoonotic in humans with gastroenteritis (61). There is, however, evidence that picobirnaviruses may actually be bacteriophage rather than eukaryote associated viruses (62), such that the virology of these viruses is currently unclear.

We identified novel fox viruses within the *Picornaviridae* belonging to three distinct genera: kobuvirus, picodicistrovirus and picornavirus. The *Picornaviridae* are a large and diverse family that include viruses associated with a variety of human diseases such as hand, foot and mouth disease, polio, myocarditis, hepatitis A virus and rhinovirus (63). All viral sequences here were most closely related to those viruses previously found in dogs. While we cannot assume that these viruses cause disease, kobuviruses have been isolated from dogs and other mammals with diarrheic symptoms (64, 65). Additionally, the fox picornaviruses found here are closely related to sapeloviruses that cause encephalitis in domestic pigs (66-68).

Finally, and of particular note, we identified rabbit haemorrhagic disease virus-2 (RHDV2) in fox faeces. RHDV was initially released (or escaped) in Australia in 1995 following testing as a biological control agent for invasive rabbits. A novel variant of the disease, RHDV2, began circulating in Australia in 2015 and is presumed to be an incursion from Europe where it first emerged in 2010 (69). RHDV2 has become the dominant strain circulating in Australia’s wild rabbits (70). The virus identified here was most closely related to RHDV2 strains found in rabbits in New South Wales, Australia in 2015-2016. It is likely, then, that Sydney foxes consume diseased rabbits and the virus is simply a gut contaminant with no active RHDV2 replication in the fox host. Although it is worth noting that antibodies against RHDV have been detected in red foxes in Germany, there was no evidence of illness or viral replication (71). Alternatively, it is possible that RHDV2 found in foxes was the result of infected fly consumption while scavenging. RHDV can be transmitted by flies after contact with diseased rabbit carcasses and remain viable for up to 9 days (72). The virus can also be excreted in fly faeces and regurgitate, which contain a sufficient number of virions to infect rabbits (72). Indeed, flies may be important vectors for pathogen transmission for scavenging predators such as foxes.

Urbanisation influences pathogen exposure and prevalence in wildlife. For example, the prevalence of parvovirus increases with proximity to urban areas in grey foxes (*Urocyon cinereoargenteus*) in the US (73), and dogs in urban areas in Brazil harbour more tick-borne pathogens than rural dogs (74). In addition, the prevalence of West Nile virus in wild birds in the US increases with proximity to urban areas and human population density (75). Here, we found the highest overall viral abundance in rural foxes while urban foxes harboured a higher diversity of viruses (Figure 2b-2c). It has previously been suggested that red foxes in highly urbanised areas experience lower exposure to canine distemper virus due to reduced movement opportunities as a result of wildlife corridors being absent in densely built-up areas (70). By comparison, exposure to canine distemper virus increased in areas with more natural habitats (76). Urban green spaces or remnant forest may therefore increase the potential for pathogen transmission due to a greater convergence of urban wildlife together with domestic animals and humans (76). This emphasises the need for targeted control of foxes in urban areas of Australia since green spaces and remnant forests have benefits associated with the conservation of native biodiversity and associated ecosystem services.

It is possible that urban living reduces fox susceptibility to viral infection by positively influencing host immunity. For example, an abundance of rich food sources would increase nutritional intake, positively influencing overall health and condition and hence resistance to viral infections (77). Kit foxes (*Vulpes macrotis*) in urban areas in California show less nutritional stress, increased body condition and improved immune function when compared to foxes in a nearby nature reserve (78). Australian lace monitors (*Varanus varius*) consuming human refuse experience improved body condition and reduced blood parasite infection compared to those that do not subsist on anthropogenic food waste (79). Foxes in urban Sydney grow larger and are heavier than foxes in rural areas (22), and there may be an advantage to consuming anthropogenic food sources for overall condition and pathogen resistance.

Across both rural and urban habitats we observed that female foxes harboured a higher abundance, and had almost twice the diversity of viruses found in male foxes (when including both vertebrate and non-vertebrate associated). While other studies looking at sex differences and immunity suggest females typically display stronger immune responses and reduced pathogen load compared to males (80), this observation could be explained by greater sociality in female compared to male foxes. That is, female foxes associate with both cubs and other ‘helper females’, whereas males are more solitary (25, 26). Greater sociality increases viral transmission opportunities, although our understanding of red fox sociality in Australia is limited (81) and males may be more likely to be involved in aggressive encounters with conspecifics than females (82). Alternatively, other biological differences, such as hormones, could also contribute to variation between male and female viromes (83).

Multiple co-occurring factors could simultaneously affect viral infection in Sydney’s foxes. Additional assessments of habitat structures, fox densities, movement behaviours and social dynamics in urban and rural areas in the Greater Sydney region will help to elucidate such factors. An obvious extension to this work is to examine fox viromes across a more comprehensive urban-rural gradient, including foxes from more isolated bush habitats. This would help us to understand differences in pathogen prevalence and transmission between isolated natural habitats and more disturbed environments, and how introduced species such as foxes contribute to disease prevalence across different ecosystems. Another useful approach could compare viral transmission dynamics in red foxes between their native and introduced ranges.

Human encroachment on wild environments and the adaptation of wild animals to urban areas continues to intensify human-wildlife interactions. The effects of urbanisation on wildlife pathogen dynamics may have unexpected consequences for human and domestic animal health. Although we cannot say definitively that the viruses identified here cause disease outbreaks or spill-over events, it is clear that foxes living in Greater Sydney carry viruses that are related to those found in domestic animals and humans. Our findings indicate that foxes may be reservoirs for viral pathogens with zoonotic potential.

## Acknowledgements

This work was funded by a grant awarded to AJRC and JLG by the Greater Sydney Local Land Services. AJRC is a recipient of the Macquarie University Research Fellowship. ECH is supported by an ARC Australian Laureate Fellowship (FL170100022).

